# Control of limb loading during active horizontal perturbations at moderate and fast trots in rats

**DOI:** 10.1101/2024.12.30.630770

**Authors:** Emanuel Andrada, Martin S. Fischer, Heiko Stark, Dirk Arnold

**Author notes:** Corresponding author: Dirk Arnold.

## Abstract

To understand how small animals cope with complex, unstructured, and unpredictable substrates, we analyzed the kinetics of rats (n = 10) moving at a fast and at a moderate trot over an unperturbed substrate and a substrate subjected to active horizontal perturbations. Perturbations were active single forwards or backwards displacements of an instrumented platform by amplitudes of 5mm or 10mm in 0.05s. Single leg ground reaction forces (SLGRF) were collected for unperturbed and perturbed locomotion (hindlimbs: 50/102, forelimbs: 45/130, respectively). When negotiating horizontal perturbations, rats displayed gait resetting (braking, accelerating) and non-resetting behaviors. Feedforward strategies differed between the fore- and hindlimbs. In circa 60% of the perturbed trials, forelimbs started the step in acceleration mode, while hindlimbs began the stance mostly in non-resetting mode (∼45%). In about 50% of all perturbed steps, the impulse provided by the SLGRF displayed a change in behavior according to the expected response to the perturbation. The remaining 50% retained the feedforward strategy. Still, most perturbed trials displayed changes in SLGRF patterns that indicated passive and active reactions to platform shifts.

Our results indicate that rats’ sensorimotor control system tunes fore- and hindlimbs differently in expectation of a perturbation. In addition, the tendon-muscle systems of the limbs are recruited to prevent leg collapse at the beginning and end of the stance. At lower speeds, spinal and/or higher center commands have enough time to re-adapt limb behavior. At higher speeds of locomotion, rats rely more on their limbs’ intrinsic stability and on feedforward control.

## Introduction

Small animals can cope with complex, unstructured, and unpredictable substrates easily by adaptively combining the intrinsic stability of their body with neuronal control ^1–3^. In the wild, small animals must negotiate natural, inherently rough terrains at high speeds. This leads to very short stance times in which to re-adapt limb load and thus posture after perturbations. Accordingly, it has been shown that animals negotiating uneven terrains preadjust limb kinematics and impedance before touch-down to reduce the need for neuronal feedback control ^3–9^. Similarly, after a sudden drop, running birds’ muscular force response is explained by intrinsic tendon-muscle properties rather than by neuronal modulation ^7,10^. Intrinsic stability plays an important role in balance during running when the touch-down (TD) of a limb occurs immediately after a perturbation. However, whether and how animals adapt leg load to negotiate perturbations arising during the stance phase has yet to be explored. Characterizing how weight-bearing limbs respond to external perturbations during locomotion may help to confirm the importance of intrinsic stability and / or to identify other neuromechanical control strategies for agile and robust locomotion.

In the present paper we analyze the kinetics of rats moving at moderate and fast trots over unperturbed and actively horizontally perturbed substrates. To obtain the necessary data, we used a novel platform for neuromechanical experiments that we termed ‘the shaker’ ^11^ which can collect ground reaction forces during active perturbations.

Measuring ground reaction forces (GRF) is a sensitive and non-invasive way to analyze the contribution of limbs to weight bearing, propulsion and stability ^12–16^. Previous work has shed light on level^17^, inclined^18^, and uneven locomotion in rats^19^. Other studies have helped to better understand differences in leg coordination between healthy rats and a) rats after cortical tract lesions^13^, b) rats after spinal cord injury^20^, and c) hemi-parkinsonian rats^14^. To the best of our knowledge, no studies exist that have analyzed horizontal substrate shifts during rat or any other small mammal locomotion.

We collected GRFs from rats moving at moderate and fast trots over unperturbed and actively horizontally perturbed platform. Backwards motions of the platform were termed caudal perturbation, and forward motions of the platform cranial perturbation (see Fig. 1 and methods for further information).

**Figure 1:**
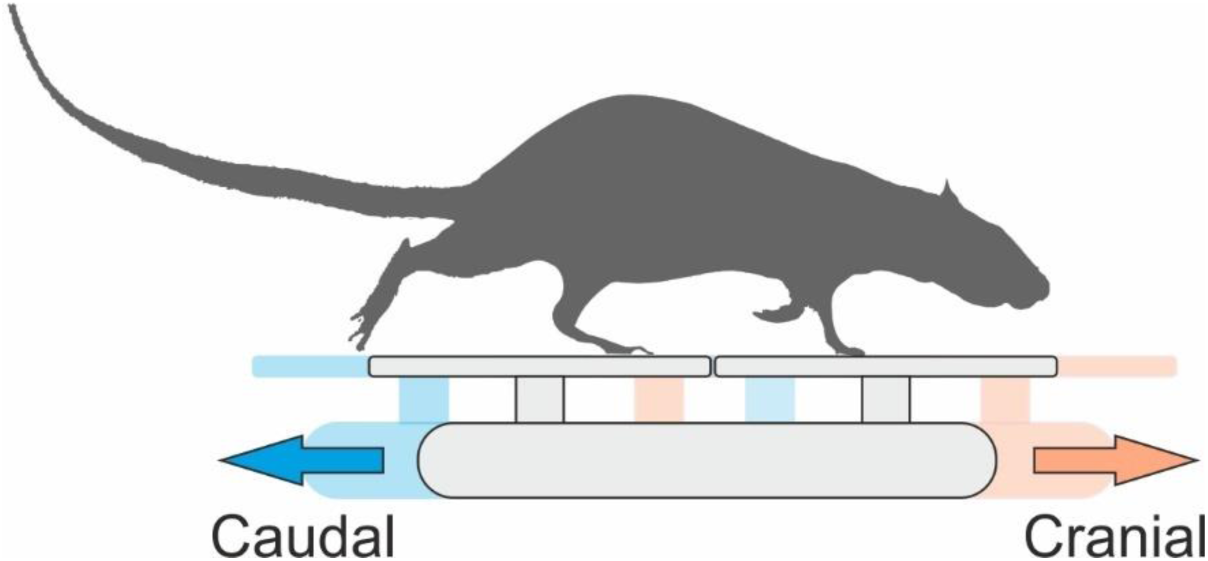
Active horizontal perturbations. The rats experienced randomly fast horizontal substrate shifts in two directions. During caudal perturbation, the platform moved backwards 5 mm (Cau5) or 10 mm (Cau10) in 50 ms. During cranial perturbation the platform moved forwards 5mm (Cra5) or 10 mm (Cra10) in 50 ms.

Since we were interested in the response to active horizontal perturbations, we compared vertical and fore-aft forces exerted during unperturbed locomotion to forces exerted in response to perturbations.

We expected that both the amplitude of the horizontal perturbation and locomotion speed would influence sensory feedback and vestibular control. We are aware that we cannot separate the effects produced by these afferent and descending signals to leg control. However, animals and humans suppress vestibular input during fast running and rely instead on spinal control modulated by type Ia sensory fibers ^21–23^. In addition, group I afferents and hip angle extension trigger the transition between stance and swing phase ^24–28^. Thus, for slower locomotion speeds and larger perturbations, we expected limb load and contact times (CT) to differ (significantly in the case of CT) from those observed in unperturbed locomotion. For the fastest trials, we expected less significant deviation from unperturbed locomotion independent of perturbation type. This last assumption is also based on the inherent stability of the highly automated quadrupedal fast trot.

## Results

We collected locomotion data from ten rats during unperturbed locomotion and from nine during horizontal perturbations (two directions: cranial and caudal, and two amplitudes: 5mm in 0.05 s and 10 mm in 0.05 s, see Fig. 1). For the hindlimbs, we collected 50 steps during unperturbed and 102 steps during perturbed locomotion. For the forelimbs, we obtained 45 steps during unperturbed and 130 steps during perturbed trials (see table 1). Rats displayed gait resetting (braking, accelerating) and non-resetting behaviors when coping with horizontal perturbations. In non-resetting trials, the shape and amplitude of the fore-aft limb forces (GRF_fa_) was like the mean curve of the GRF_fa_ during unperturbed locomotion. Few rats stopped the first time they experienced a perturbation (not included in our results). After one or two perturbed trials, the rats knew that a perturbation might occur when they stepped on the shaker. Thus, in the following trials the rats tended to accelerate to cross the perturbing platform. Interestingly, many of them suppressed that accelerating behavior after a couple of trials and just performed a non-resetting trot program independent of the perturbation.

**Table 1.** Animals, weights, and collected steps.

| Rat name | Weight [g] | Steps unperturbed |  | Steps cra5 |  |  |  | Steps cra10 |  |  |  | Steps cau5 |  |  |  | Steps cau10 |  |  |  |
| --- | --- | --- | --- | --- | --- | --- | --- | --- | --- | --- | --- | --- | --- | --- | --- | --- | --- | --- | --- |
|  |  |  |  | E |  | L |  | E |  | L |  | E |  | L |  | E |  | L |  |
|  |  | F | H | F | H | F | H | F | H | F | H | F | H | F | H | F | H | F | H |
| Right | 400 | 11 | 7 | 6 | 4 | 6 | 1 | 1 | 1 | 4 | 2 | 2 | 3 | 3 | 2 | 4 | 2 | 5 | 2 |
| Left | 290 | 4 | 5 | 3 | 5 | 4 | 2 | 1 | 6 | 6 | 2 | 2 | 7 | 2 | 2 | 1 | 4 | 2 | 1 |
| Black 1 | 300 | 4 | 3 | 3 | 0 | 0 | 0 | 2 | 2 | 2 | 0 | 0 | 1 | 2 | 0 | 0 | 1 | 2 | 0 |
| Black 2 | 295 | 4 | 3 | 1 | 1 | 1 | 1 | 3 | 2 | 1 | 2 | 1 | 1 | 3 | 2 | 2 | 2 | 0 | 0 |
| Black 3 | 303 | 1 | 3 | 1 | 1 | 1 | 0 | 1 | 1 | 1 | 0 | 2 | 1 | 1 | 1 | 1 | 2 | 0 | 0 |
| Black 4 | 298 | 4 | 4 | 0 | 0 | 3 | 1 | 1 | 2 | 0 | 0 | 2 | 1 | 1 | 0 | 2 | 1 | 2 | 0 |
| Red 1 | 296 | 5 | 5 | 4 | 7 | 3 | 0 | 3 | 2 | 1 | 0 | 2 | 1 | 2 | 1 | 1 | 1 | 1 | 0 |
| Red 2 | 301 | 7 | 6 | 4 | 0 | 1 | 0 | 0 | 0 | 0 | 0 | 0 | 0 | 1 | 0 | 1 | 2 | 3 | 1 |
| Red 3 | 305 | 3 | 3 | 0 | 0 | 0 | 0 | 0 | 0 | 0 | 0 | 0 | 0 | 1 | 0 | 0 | 0 | 0 | 0 |
| Red 4 | 290 | 7 | 6 | 1 | 2 | 4 | 3 | 2 | 3 | 2 | 1 | 3 | 2 | 3 | 2 | 1 | 1 | 1 | 0 |

E: Early stance, L: Late stance, H: Hindlimb, F: Forelimb.

### Perturbed vs. unperturbed locomotion

#### Hindlimbs

During unperturbed trot locomotion, the contact time (CT) of the hindlimbs was on average 0.164 ± 0.04 s and 0.09 ± 0.01 s for the moderate and fast trot groups, respectively (Table 2). Significant differences were found between the trot groups (moderate vs fast trot, for all perturbation types p < 0.05). No significant differences were found between perturbed and unperturbed locomotion, independent of the timepoint of the onset of the perturbation (see tables 2 and S3).

During unperturbed locomotion, the maximum value and the timepoint of occurrence of the vertical component of GRF (GRF_v_) were on average 0.73 ± 0.1 of body weight (BW=force/(mass*gravity) at 33 ± 8 % of the stance for moderate trot, and 0.96 ± 0.12 BW at 37 ± 5 % of the stance for fast trot. The maximum value of GRF_v_ differed significantly (p < 0.01) between trot groups, but the timepoint did not. GRF_fa_ exhibited a mean minimum negative value of −0.09 ± 0.03 BW at 12 ± 5 % of the stance for moderate trot, and a mean minimum value of −0.08 ± 0.03 BW at 5 ± 3 % of the stance for fast trot. The timepoint of the minimum value differed significantly (p < 0.003) between trot groups, but the amplitude did not. During fast trot the maximum positive GRF_fa_ was larger on average (fast trot: 0.22 ± 0.1 BW vs moderate trot: 0.15 ± 0.06 BW, p > 0.05) and was reached significantly earlier than during moderate trot (fast trot: 53 ± 4 % vs moderate trot: 65 ± 8 %, p = 0.0002).

For all early stance perturbations (ESP) regardless of perturbation amplitude and direction, the maximal peak value of the GRF_v_ was larger on average for the fast trot. However, the difference was only significant for cranial translations of 5 mm in 50 ms (Cra5) (Cra5-fast vs Cra5-moderate, p = 0.046). The peak GRF_v_ value for fast trot occurred later during stance than in moderate trot (p > 0.05). No significant differences were found between unperturbed and perturbed or between perturbed trials (5 mm vs. 10 mm and cranial vs. caudal) in the same trot speed group (moderate or fast). Neither speed nor perturbation type significantly influenced the min and max values or timepoint of the GRF_fa_. For more information see Supplementary Tables.

However, fast trotting rats exhibited earlier and lower negative GRF_fa_ peak values and later (with the exception of cranial perturbations) and larger maximal positive GRF_fa_ peak values on average than rats trotting moderately. Fast locomotion induced on average an earlier transition from braking to accelerating GRF_fa_. A similar earlier transition was observed between moderate and fast unperturbed locomotion.

For late stance perturbations (LSP) of fast trots the maximum peak value of the GRF_v_ was larger, on average (though only significantly larger in the case of Cau5-fast vs Cau5-moderate, p = 0.02) and occurred later than during perturbed moderate trots. Maximum GRF_fa_ values and GRF_fa_ timepoints obtained during perturbed locomotion were not significantly different from those obtained during unperturbed trials. However, maximum GRF_fa_ values obtained during caudal perturbations were lower on average than those obtained during cranial ones. For more information see Supplementary Tables.

The combination of the impulse gap (IG) and the square of IG relates the kinetic behavior of a perturbed rat limb to the average kinetic behavior observed during non-perturbed locomotion (see methods). Three related behaviors are possible: acceleration, braking and non-resetting. We analyzed IG and IG^2^ combinations for the whole stance, and for the stance periods located before and after the onset of the perturbation. IG and IG^2^ values for the hindlimb’s whole stance phase indicated that rats prefer two strategies when negotiating caudal perturbations: a) braking and b) a non-resetting behavior. Perturbed hindlimbs braked in about 40% of ESP trials and in 35% of LSP trials and displayed non-resetting behavior in roughly 50% of both the ESP and LSP trials (Figs. 2A & 2F, Table S10). When mapping IG and IG^2^ before and after the perturbation (plot embedded in Figs. 2A and 2F), we found that 42% of the perturbed steps started in acceleration mode while about 45% started in non-resetting mode. Most of the steps that started in acceleration mode (∼57 %) turned non-resetting after the perturbation. The rest remained accelerative (pure acceleration) or braked after the perturbation, in equal proportions (about 21% each). From the steps that started in non-resetting mode, circa 50% remained so, while the other half slowed down. With only one exception, the few steps that began in braking mode remained in that mode (see Fig. 2 A).

**Figure 2.**
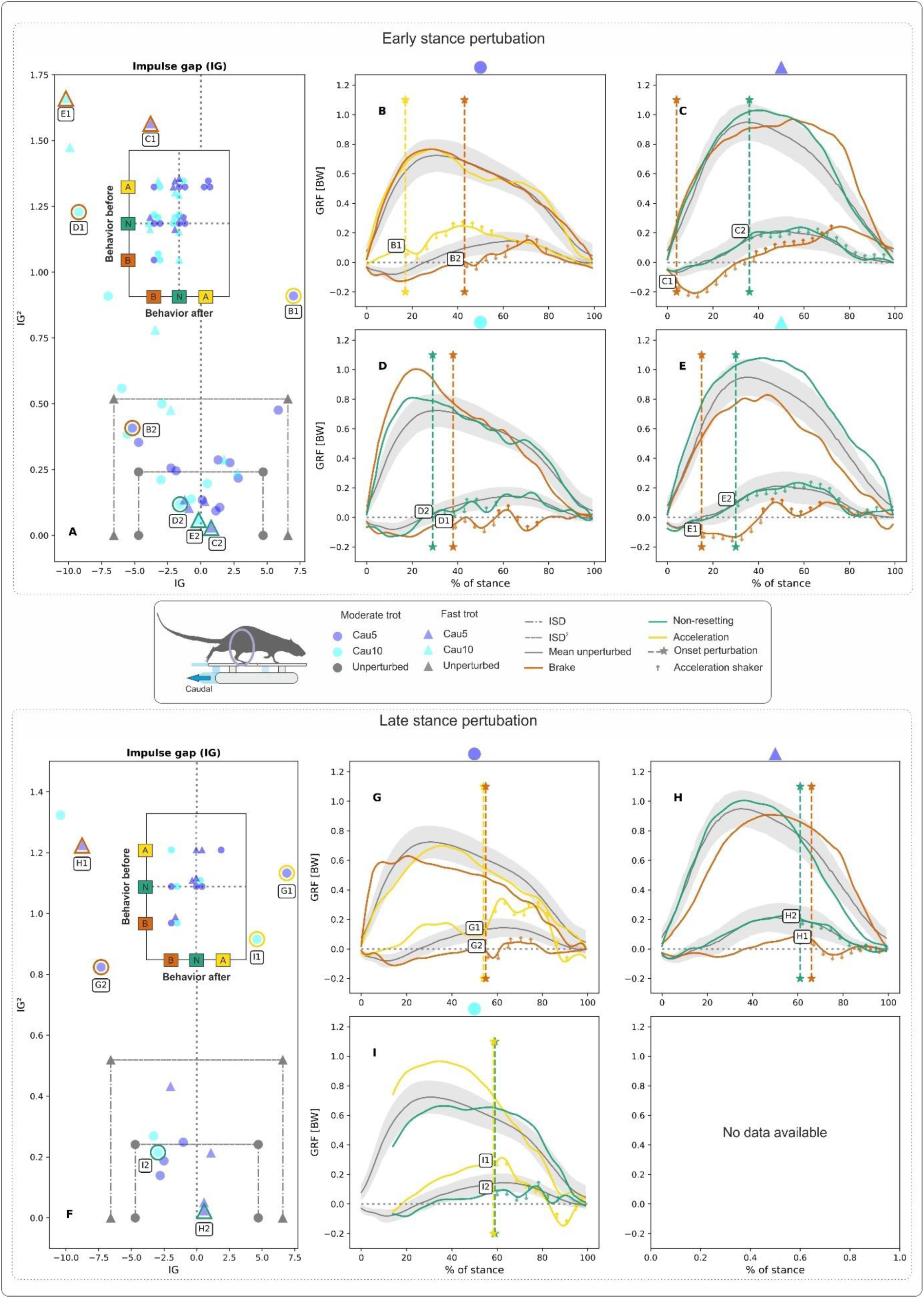
Influence of backward translations of the shaker (here termed caudal perturbations) on hind limb kinetics during rat locomotion. Perturbations occurred in the early (A-B-C-D-E) and in the late stance (F-G-H-I) phases at moderate and fast trots. Deviations in the patterns of the ground reaction forces (GRF) with respect to those obtained during unperturbed locomotion were captured in the form of the impulse gap (IG) (see A & F). IG provides a measure of the shift between perturbed and unperturbed fore-aft GRF during stance. Quadratic IG values in the vertical axis of figs. A & F provide a measure of the amplitude of the oscillation of the perturbed fore-aft GRF around the unperturbed ones. IG and IG^2^ values relate perturbed locomotion to unperturbed locomotion. Positive values for IG indicate that the rats accelerated, negative ones that the rats decelerated. For non-resetting behavior, both IG and IG^2^ values must lie in the area formed by the two vertical and the horizontal dot-dashed lines with markers (circle for moderate and triangle for fast trot). The vertical lines represent the sum of the standard deviation (SD) of the GRFs from unperturbed trials, here referred to as the impulse margin (ISD), while the horizontal line represents their quadratic value, ISD^2^ (A & F). The map of behaviors before and after the perturbation is shown in the plots embedded in A & F. The diagonal from bottom left to top right displays pure behaviors. B-C-D-E-G-H-I display selected examples of GRF from perturbed trials (marked in A & F with circles or triangles, subplot letter and curve number) superimposed on the average profiles ± one SD of GRFs from unperturbed trials. Vertical dashed lines with stars in B-C-D-E-G-H-I denote the onset of the perturbation. Note that the shaker’s horizontal shifts have acceleration and deceleration ramps. These ramps, with a duration of circa 25 ms each, are marked with arrows in the fore-aft GRF curves of the perturbed trials in B-C-D-E-G-H-I. Cau5: 5mm backward shifts of the platform, and Cau10: 10 mm backward shifts of the platform, both in 0.05 s).

Figures B-C-D-E-G-H-I display selected examples of GRFs from perturbed trials superimposed to the average profiles ± SD of GRFs from unperturbed trials. In figures B-C-D-E, responses to ESP can be seen. After the onset of the perturbation, the curves of the GRF_fa_ generally display a dip followed by an increase in the fore-aft forces. At a moderate trot, the dip leads to oscillations that continue for several periods (e.g., Fig. 2D, curves D1 and D2). At a fast trot, on the other hand, GRF_fa_ oscillations are significantly lower, especially in trials in which non-resetting behavior was observed (e.g., Figs. 2 C-E, curves C2 and E2). Responses to LSP display similar patterns to those found in responses to ESP (Figs. 2 B-C-D).

When negotiating substrate shifts in a cranial direction, rat hindlimbs displayed more pure accelerative behavior than when negotiating substrate shifts in a caudal direction (see quadrant A-A in plot embedded in Fig. 3 A). However, in 60% of the trials, rats coped with perturbations without engaging pure accelerating or braking strategies. Pure braking was seldomly observed (quadrant B-B in plot embedded in Fig. 3 A-D). Similarly, accelerative responses after the onset of the perturbation in trials that started in braking or non-resetting modes were infrequent (B-A quadrant or N-A region in plot embedded in Fig. 3 A-D).

**Figure 3.**
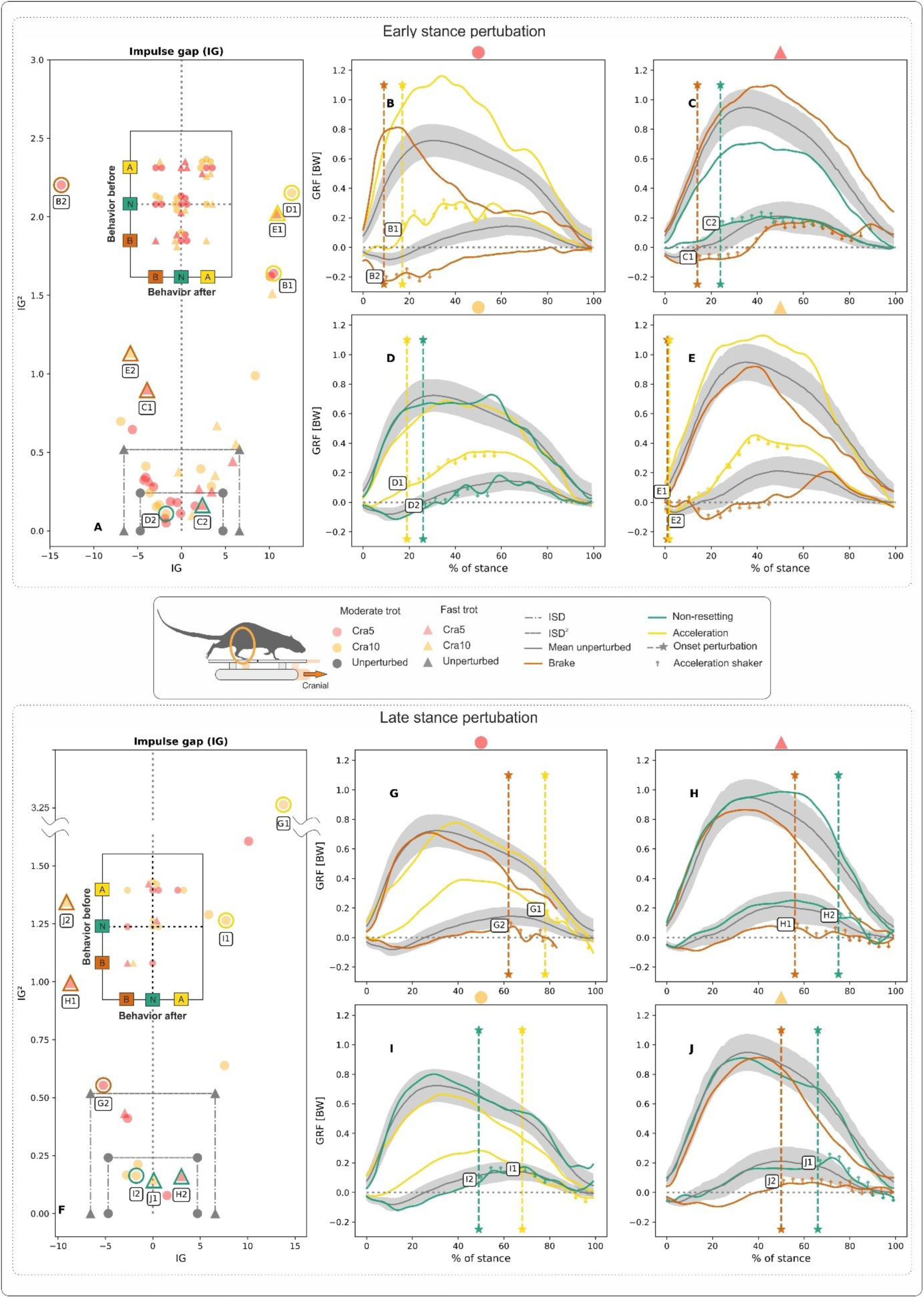
Influence of forward translations of the shaker (here termed cranial perturbations) on hind limb kinetics during rat locomotion. Perturbations occurred in the early (A-B-C-D-E) and in the late stance (F-G-H-I) phases at moderate and fast trots. Deviations in the patterns of the ground reaction forces (GRF) with respect to those obtained during unperturbed locomotion were captured in the form of the impulse gap (IG) (see A & F). The map of behaviors before and after the perturbation is shown in the plots embedded in A & F. B-C-D-E-G-H-I display selected examples of GRF from perturbed trials superimposed on the average profiles ± one SD of GRFs from unperturbed trials. Cra5: 5mm forward shifts of the platform, and Cra10: 10 mm forward shifts of the platform, both in 0.05 s). See Fig. 2 for further information.

Cranial early stance perturbations did not significantly influence the shape of the GRF_v_. However, Fig.3 D displays one trial in which non-resetting behavior was observed. In this example, a bump can be seen in the GRF_v_ at the beginning of the decelerative part of the perturbation (Fig.3 D, curve D2). In trials in which non-resetting was observed, the GRF_fa_ displayed a bump after a cranial late stance perturbation (see e.g., Figs. 3H and 3J, curves H2, and J1, respectively), while the GRF_v_ did not obviously show late stance perturbation-related changes. As observed for caudal perturbations, responses in GRF_fa_ after cranial late stance perturbations when rats were moving at a fast trot seem to be more dampened.

#### Forelimbs

During unperturbed trots, the contact time of the forelimbs was 0.166 ± 0.04 s and 0.09 ± 0.01 s (p < 0.0001) on average for moderate and fast trot groups, respectively.

Significant differences (p < 0.05) were also observed in perturbed trials between the contact times of the two gait groups (moderate vs fast trot. However, within trot groups (moderate or fast), our results showed no significant differences in contact times between perturbed and unperturbed locomotion, regardless of the timepoint of the onset of the perturbation. For more information see Table S2 and other Supplementary Tables.

During unperturbed locomotion, the maximal value of the vertical component of GRF (GRF_v_) and its point of occurrence were on average 0.7 ± 0.1 BW at 59 ± 12 % of the stance and 0.98± 0.1 BW at 54 ± 8 % of the stance for moderate and fast trot groups, respectively. The maximal value of the GRF differed significantly between trot groups (p < 0.001). The fore-aft component of the GRF (GRF_fa_) exhibited a mean minimal negative value of −0.1 ± 0.04 BW at 24 ± 9 % of the stance for moderate trot and a mean minimal value of −0.085 ± 0.03 BW at 26± 14% of the stance for fast trot. Differences between trot groups were not significant. The maximal positive value of GRF_fa_ was roughly the same for both trot groups 0.093 ± 0.03 BW, but during fast trot the maximal value was reached somewhat earlier (76 ± 7 % vs 80 ± 5 %) in the stance phase. Differences between trot groups were not significant.

For early stance perturbations, the fast and moderate trot groups formed two separated data clusters as regards the GRF_v_ (see Table S7). Regardless of the perturbation type, the maximal peak GRF_v_ values were significantly larger (p < 0.05) and occurred earlier during stance (p > 0.05) during the fast trot than during the moderate trot. However, when comparing unperturbed with perturbed trials or when comparing the different perturbation scenarios, no significant differences were found within trot speed groups (moderate or fast). During perturbed locomotion, minimal and maximal peak GRF_fa_ values and timepoints did not differ significantly between trot speed groups (moderate vs. fast). Nor did they differ significantly between the different perturbation scenarios or between perturbed and unperturbed locomotion. For more information see Supplementary Tables.

For late stance perturbations, maximum GRF_v_ values differed significantly between the two trot groups (p < 0.05). As regards timing, peak maximal force was reached earlier during the fast trot, but results did not differ significantly between the trot groups (p > 0.05). In perturbated trials, maximum GRF_fa_ values were larger on average when perturbations occurred late in the stance phase. However, neither the maximal values nor the timepoint of their occurrence differed significantly from unperturbed locomotion or between the different perturbation scenarios (p > 0.05). For more information see Supplementary Tables. Some differences emerge between the forelimbs and the hindlimbs in their response to caudal and cranial perturbations.

When the stance phase is analyzed as a whole in trials with caudal shifts, IG and IG^2^ reveal that forelimbs accelerated in 46% and braked in 31% of the runs. Only about 23% of the perturbed steps were purely non-resetting.

Before a caudal shift, most of the steps started in acceleration mode (∼62% of all ESP and LSP trials). Circa 57% of ESP and 40% of LSP steps remained purely accelerative, although the logical response to a caudal shift would be to brake (see quadrant A-A in the embedded plots in figures 4A and 4F). However, the percentage of trials in which the forelimbs abandoned the acceleration mode after the onset of the perturbation increased significantly from ESP to LSP (from 15% to almost 50%, respectively).

**Figure 4.**
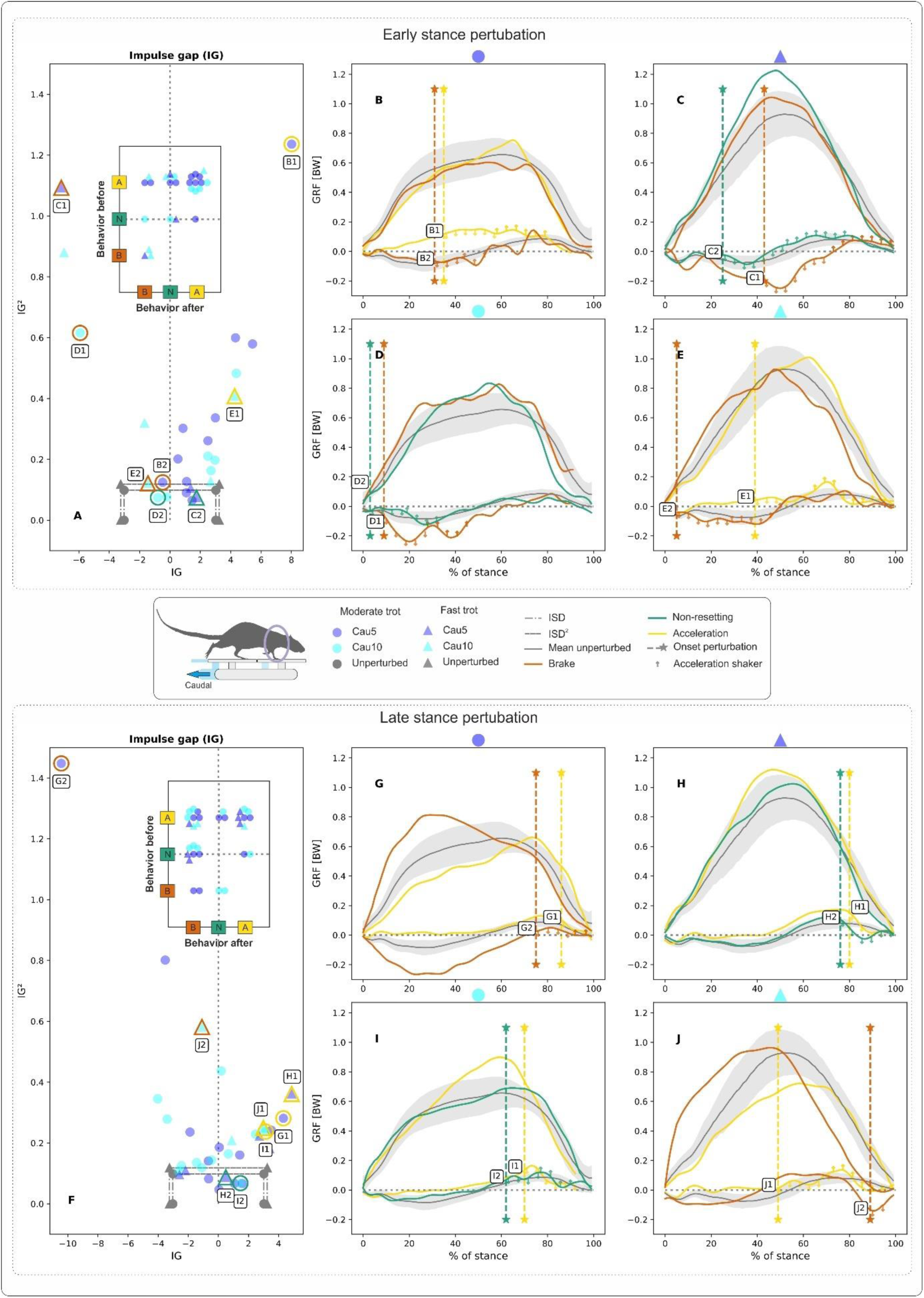
Influence of backward translations of the shaker (here termed caudal perturbations)on forelimb kinetics during rat locomotion. Perturbations occurred in the early (A-B-C-D-E) and in the late stance (F-G-H-I) phases when rats were moving at moderate and fast trots. Deviations in the patterns of the ground reaction forces (GRF) with respect to those obtained during unperturbed locomotion were captured in the form of the impulse gap (IG) (see A & F). The map of behaviors before and after the perturbation is shown in the plots embedded in A & F. B-C-D-E-G-H-I display selected examples of GRF from perturbed trials superimposed on the average profiles ± one SD of GRFs from unperturbed trials. Cau5: 5mm backward shifts of the platform, and Cau10: 10 mm backwards shifts of the platform, both in 0.05 s). See Fig. 2 for further information.

Purely non-resetting steps were sparse. In addition, 60% of the steps that started as non-resetting shifted into braking mode after the perturbation. This was more marked for the LSP trials (see region N-B in the embedded plot in Fig. 4F and a single trial in Fig. 4H, curve H2). As already mentioned for the hindlimbs, most of the (few) braking steps measured did not change mode after the perturbation (see e.g., region B-B in the embedded plot in Fig. 4A).

Forelimb kinetic responses to cranial shifts are presented in Fig 5. IG and IG^2^ values for the stance phase as a whole indicated that rats’ forelimbs accelerated in ∼50% of perturbed steps, while they braked or showed non-resetting behavior in circa ∼25% of the perturbed steps.

**Figure 5.**
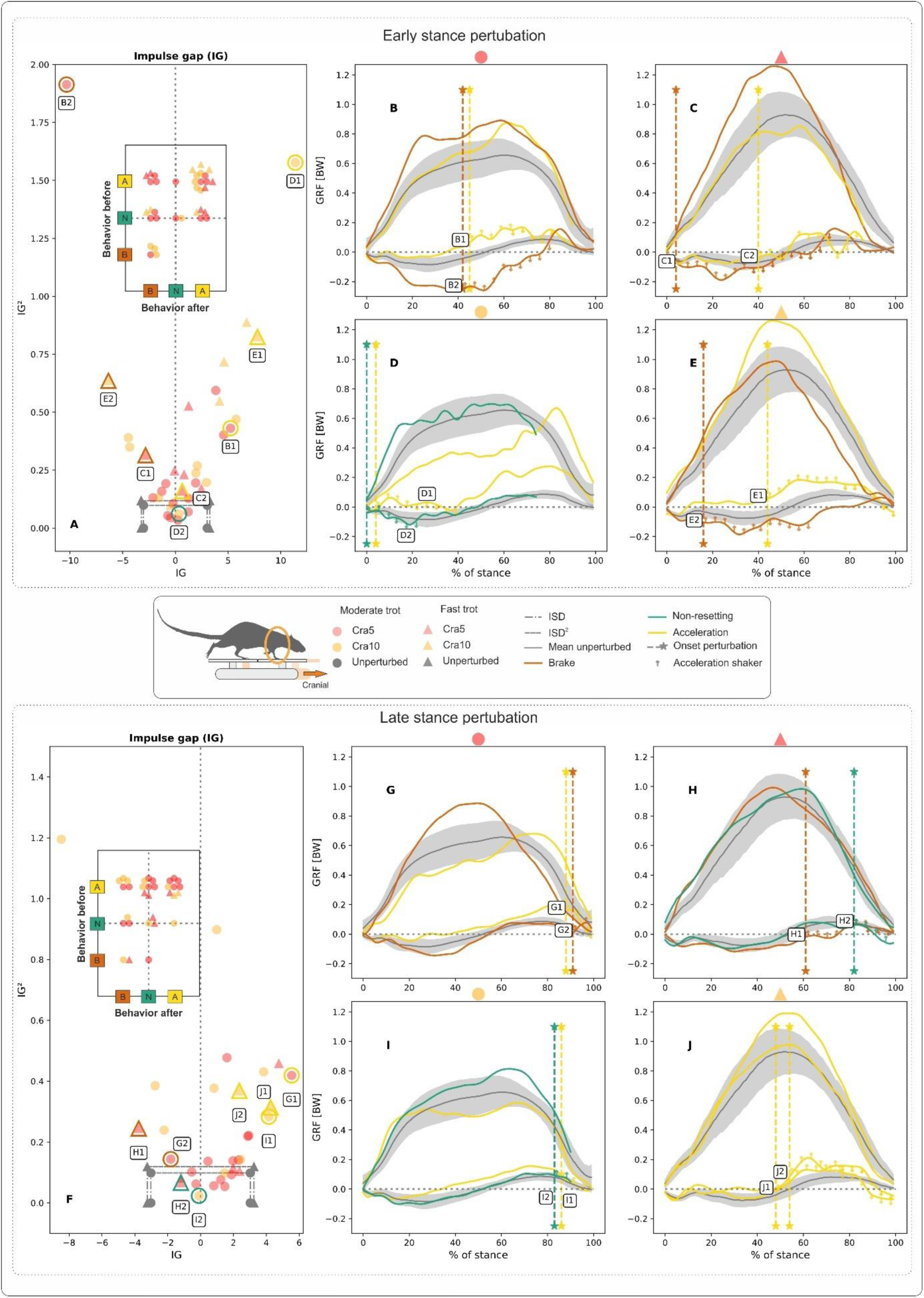
Influence of forward translations of the shaker (here termed cranial perturbations) on forelimb kinetics during rat locomotion. Perturbations occurred in the early (A-B-C-D-E) and in the late stance (F-G-H-I) phases at moderate and fast trots. Deviations in the patterns of the ground reaction forces (GRF) with respect to those obtained during unperturbed locomotion were captured in the form of the impulse gap (IG) (see A & F). The map of behaviors before and after the perturbation is shown in the plots embedded in A & F. B-C-D-E-G-H-I display selected examples of GRF from perturbed trials superimposed on the average profiles ± one SD of GRFs from unperturbed trials. Cra5: 5mm forward shifts of the platform, and Cra10: 10 mm forward shifts of the platform, both in 0.05 s. See Fig. 2 for further information.

The mapping of behaviors observed before and after a perturbation resulted in roughly similar patterns for both ESP and LSP (cp., embedded plots in Figs. 5A & 5F). In ∼60% of the collected steps, rats started by accelerating with their forelimbs. After an ESP, the accelerating forelimb remained in acceleration mode in almost 70% of cases. A shift to braking/non-resetting modes happened only in 25% / 5% of the trials, respectively. After an LSP, forelimbs remained in acceleration mode in 41% of cases and switched to non-resetting or braking modes in 36% and 23% of the trials, respectively.

Only about 30% of the trials that started out as non-resetting remained purely non-resetting. Of the remaining 70%, ∼40% shifted to braking mode, while the other ∼30% shifted to acceleration mode after the onset of the perturbation.

In terms of GRF_v_ and GRF_fa_ patterns, forelimb responses after a caudal/cranial perturbation did not differ from those identified for the hindlimbs. In general, GRF_v_ did not exhibit clear changes after either caudal or cranial perturbations. However, rats that were perturbed at a moderate trot and were in acceleration mode displayed a bump in GRF_v_ at the end of the stand (see Figs. 4B, 4D, 5B-5B-5E). Having said this, the same pattern is sometimes also to be observed in purely non-resetting trials (see Fig. 5I).

A caudal shift often induced a deceleration dip in the GRF_fa,_, while a cranial shift induced an acceleration bump/ramp. These dips or bumps/ramps sometimes appeared immediately after the perturbation (e.g., Figs. 4C & 4D, curves C1 & D1, Figs. 5D & 5G, curves D1 & G2) and in other cases with a delay (e.g., Figs. 4B & 4D, curves B2 & D2, Figs. 5B-5H, curves B2 & H1). Sometimes, dips or bumps led to oscillations, particularly during a moderate trot (see Figs., 4B & 4D, curves B2 & D1, Figs. 5B & 5C, curves B1 and C2). Finally, Fig. 4B (curve B1) shows a trial which displays no clear response to a caudal perturbation. The complete set of statistical tests and their results can be found under following link: http://rat.stark-jena.de.

## Discussion

To infer the control strategies implemented by a dynamic system, it is useful to identify the ways in which the system responds to external perturbations. We aimed to establish how weight-bearing limbs respond to horizontal shift perturbations during locomotion. We hoped to shed light on the trade-off between the feedforward and feedback strategies employed during stance that ensure agile and robust locomotion. Feedforward strategies are generated from higher brain centers before the limb makes contact with the ground and are based on sensory perceptions and internal models. In contrast, feedback modulation of motor output occurs during limb contact and is said to be coordinated via short-latency reflex feedback to spinal circuits ^8,29–31^.

To analyze this trade-off, we compared rat single leg GRFs during unperturbed locomotion with rat single leg GRFs collected during trials in which rats had to negotiate rapid horizontal substrate shifts.

On the basis of previous findings ^24–28^, we hypothesized that, at lower trot speeds in particular, differences in limb load mediated by Ia afferents might induce differences in contact times compared to unperturbed locomotion. However, we found no significant differences between perturbed and unperturbed locomotion in either speed group. Similarly, statistical methods based on analysis of variance found minimal or no differences in the several control points we proposed to analyze perturbation-related changes to the shape of the GRFs (see methods). At the same time, upon visual inspection of the shape of the GRF from perturbed trials, clear differences to the mean curves of unperturbed runs are apparent.

Using the impulse gap (IG, see methods), we were able to categorize limb responses to perturbations, regardless of amplitude, direction and timepoint of the perturbation, and regardless of trot speed, into three main behaviors: 1) braking, 2) acceleration, and 3) non-resetting.

Rats did not always display the same behavior to negotiate the same horizontal perturbation. The first time they experienced a perturbation, they usually stopped (not included in our statistical results). Once they realized that the platform may move, they exhibited resetting behaviors (slowed down or speeded up) to cross the perturbing platform. Interestingly, after a while, rats suppressed the accelerating behavior and also performed non-resetting ^32^ trot programs during perturbed trials. Thus, our results include a “learning” effect on the trade-off between feedforward and feedback control.

### Feedforward Strategies

Feedforward strategies are planned behaviors (acceleration, braking, non-resetting) in advance of an expected perturbation. Higher centers such as the cerebellum generate goal-directed movements based on sensory information and internal models before the leg performs a step on the shaker^33–35^.

Feedforward strategies differed between fore- and hindlimbs when rats faced an expected perturbation. In circa 60% of the trials, forelimbs started the step in acceleration mode, while hindlimbs began the stance phase mostly in non-resetting mode (∼45%). Our results for unperturbed locomotion showed that the standard deviation (SD) for the forelimbs was significantly narrower than that computed for the hindlimbs (for both speed categories p< 0.0001, cp. e.g., Fig. 2B with Fig. 4B). This phenomenon makes it more difficult for the IG and IG^2^ values to fit inside the non-resetting region for the forelimbs than for the hindlimbs. However, a narrow SD may indicate tighter and more precise loading control in the forelimbs, which might be powered by their ability to manipulate objects. If so, our results indicate that the forelimbs might be more likely to be tuned in expectation of a perturbation than the hindlimbs.

### Pure behaviors

We termed “pure” behaviors those behaviors which did not change in the face of a perturbation, regardless of its type or the timepoint of its onset. Pure behaviors seem to be governed by strong feedforward strategies.

Pure braking steps occurred seldom. For both fore- and hindlimbs, they were observed in less than 9% of all perturbed runs. This may be connected to the fact that, as outlined above, braking was uncommon as a feedforward strategy. Pure accelerating steps were the most usual control strategy for the forelimbs, but little observed for the hindlimbs (about 41% and 15% of all perturbed trials, respectively). Note that on average, rat forelimbs braked until about 55%-60% of the stance phase during unperturbed locomotion (see Figs 4 or 5). Thus, an accelerating impulse at the beginning of the stance configures a completely different motor control set-up in which retractor muscles in the most proximal joints must be recruited from the beginning of the stance phase.

The occurrence of pure accelerating steps decreased for LSP in both hind- and forelimbs. The most obvious explanation for this would seem to be the need to avoid excessive limb retraction in order to prevent limb collapse, especially when the platform is moving backwards.

As with the feedforward strategies, pure non-resetting steps were more often observed in the hindlimbs than the forelimbs. While one in every four perturbed hindlimb steps was pure non-resetting, only 5% of perturbed forelimb steps fitted into this category. Moreover, the frequency of occurrence of pure non-resetting behavior in the hindlimb remained similar regardless of perturbation type or timepoint of onset.

### Behavioral changes induced by perturbations

In more than 50% of perturbed trials, both fore- and hindlimb behaviors changed after the onset of the perturbation. Hindlimb behaviors changed in about 51% / 56% of the trials after caudal / cranial perturbations, respectively, while forelimb behaviors changed in 55%/ 51% of the trials. The changes observed corresponded in the main to those expected to counteract the motion of the shaker (caudal perturbation: braking, cranial perturbation: acceleration). After perturbation, hindlimbs converged more frequently towards non-resetting mode, while forelimbs tended more towards braking or accelerating behaviors. Interestingly, while acceleration mode often shifted into braking mode, shifts from braking to acceleration were extremely rare. This finding indicates that braking feedforward strategies are very attractive, so the limb usually remains in braking mode for most of the stand. Having said this, neither this finding nor the fact that in circa 50% of trials limb behavior did not change after a perturbation mean that the rat limb did not react to the horizontal shifts (see next section).

### Short and long latency reactions in fore-aft forces

Even when limb behavior did not change after a perturbation, the GRF_f-a_ often evidenced reactions that deviated clearly from the patterns depicted during unperturbed locomotion. After cranial perturbations, GRF_f-a_ exhibited acceleration bumps (e.g., Fig. 3J, curve J1) or ascending ramps (e.g., Fig. 3C & 3E, curves C1 & E1), while dips (e.g., Fig. 4H, curve H2) or descending ramps (e.g., Fig. 2I, curve I1) were observed after caudal perturbations. Because it was impossible to prevent dips and bumps being introduced via compensation of the forces created during platform motions (see methods), we carefully checked every perturbed step analyzed and can confirm that the amplitude of the bumps and dips we discuss here are two or three dimensions larger than the error we observed due to compensation and synchronization (see Fig.7).

The acceleration bumps resisted the cranial displacement of the shaker and prevented an uncontrolled protraction/flexion of the limb. The rapid responses observed indicate that cranial perturbations likely stretched the extensor limb muscles (e.g., triceps in the forelimbs and gluteus in the hindlimbs), triggering, via muscle spindles and fast myelinated Ia afferents, monosynaptic reflexes ^36,37^. In some cases, bumps occurred right after the perturbation (see e.g., Fig. 5B, curve B1, Fig. 3J, curve J1). This pattern seems at first glance to correlate with an expected reaction to the perturbation, but the reaction time was arguably too fast even for reflex responses. At a conduction velocity of about 45 ms^-1^ ^38^, the time taken for the stimuli to reach spinal centers and return to the limbs would be under 2 ms. Muscle contraction times in the rat, however, are longer than 10 ms ^39^. Thus, sensorimotor control is not likely to explain the force changes in question. However, if muscles worked isometrically, which is a necessary condition for the pantograph mechanism in quadruped limbs^40^, passive structures such as tendons might help to explain this rapid increase in limb loading. Passive force compensation has, for example, been observed in guinea fowl when negotiating sudden drops ^7,41^.

Caudal perturbations induced in most cases a dip in the GRF_fa_ of both fore and hind limbs that opposed the retraction of the limb. However, when perturbations occurred early in the stance phase, this reaction occurred later in the forelimbs than after cranial perturbations (e.g., Fig 4B, curve B2). In other words, the forelimbs retracted for several milliseconds due to the action of the active platform without this reaction being counteracted. One explanation for this finding is the notion that muscle force depends on it state (strain, strain rate, strain history and loading), (see, for example,^10,42^). On the other hand, the finding could also indicate a difference in sensitivity tuning between muscle spindles in the primarily silent flexor muscles and the active extensor muscles. Note that the γ-motoneurons allow spindle response properties to be tuned ^36^. There is experimental evidence to show that γ-efferent commands covary with joint angles during movements^37,43,44^, perhaps to ensure optimal work between muscle extensor/flexor groups ^45^. Interestingly, just before TO, the dips appear in the forelimbs immediately after a caudal perturbation (Fig. 4H, curves H1 & H2). In this part of the stance, the flexor *M. biceps brachii* is activated to prepare for the swing phase and the tendons may rapidly work against the hyperextension of the elbow.

Ascending or descending ramps with significant force increments appeared circa 10ms to 25ms after perturbation onset (see e.g., Fig. 2E, curve E1, Fig. 3C, curve C1, Fig. 5B, B1). Note that in our experiments the rat moved with the platform, the perturbation was not limited exclusively to the limbs. Thus, due to the short stimuli distances in rats, spinal control and/or higher centers like the vestibular system might have been involved the observed responses. Motor commands from higher centers are not only transmitted to the leg’s central pattern generators (CPG) via descending spinal pathways^8,27,29,31,46,47^, the lateral vestibulospinal tract has direct connections to flexor and extensor motoneurons at the knee and the ankle joints. Thus it can modulate muscle activity and in some cases can affect the timing of the rhythm^48^.

### Influence of trot speed

In our experiments, the rats trotted at self-selected speeds. To reduce the effect of speed on the results, we separated the trials into moderate and fast trots (see methods). As stated above, we found significant differences in CT and in the shape of the GRF between moderate and fast locomotion within each group (e.g., non-perturbed_moderate vs. non-perturbed_fast). However, differences were not significant when unperturbed locomotion was compared with perturbed locomotion at the same speed category (e.g., non-perturbed_moderate vs. Cra10_moderate).

At moderate (slower) trot speeds, the response to the horizontal shift of the shaker led frequently to oscillations in the GRF_fa_. Two possibilities can be considered here. A) The muscles were recruited discretely to negotiate the perturbation, or b) muscle activation became unstable. It is well known that feedback control systems involving muscles as sensor and actuator units can lead to oscillating responses if negative feedback (e.g., inhibitory signals from Renshaw cells, tendon organs, contralateral limbs) experiences a delay ^37,49^.

At a fast trot, we found that oscillations in the responses to a perturbation were significantly smaller (see Fig., 2C, Fig. 2E & Fig. 4C, curves C2, E2, and C2 respectively). Similarly, “dampened” responses were observed during pure accelerative steps (see, e.g., Fig. 3D, curve D1, Fig 4B, curve B1). To enable organisms to move faster, higher centers increase both α and γ activity. Higher γ activation levels in turn reduce the sensitivity of the spindles to muscle stretch ^36^. In terms of limb size and limb speed, rat joints lie in a viscous overdamped region at a fast trot ^50^. In this region viscous effects dissipate most energy in the joints and perturbations do not cause the limb to oscillate. As locomotion speed increases, feedback control gain decreases and rats rely more on the intrinsic stability of their body plan and feedforward control.

## Conclusion

Since quadrupedal mammals maintain ground contact with two legs during the trot, they can balance out horizontal shifts more easily than bipeds. In line with this, our experiments revealed that no single response to a horizontal shift exists. In rats, reactions to perturbation seem to result from a complex combination of expectation and experience, as well as the intrinsic stability of the body. With no experience, rats usually stopped after a horizontal perturbation. With experience, rats learned to include feedforward and feedback control, speeding up to cross the perturbing platform. With even more experience, rats often suppressed the accelerating behavior and performed a non-resetting^32^ trot program during perturbed trials. Interestingly, reactions to perturbations differed between the fore- and hindlimbs. The loading of the forelimbs seems to be more strictly controlled than in the hindlimbs. During stance, intrinsic stability and feedback control tune limb-loading according to perturbation type and timepoint of onset, feedforward strategy and experience. When the perturbation occurred at the beginning or end of the stance, the passive properties of tendon-muscle system of the limb prevented leg collapse. In between, spinal and/or non-spinal centers had enough time to readjust limb behavior at moderate speeds. At higher speeds, our results indicate that rats rely more on the inherent stability of their limbs and on feedforward control.

## Material & Methods

### The shaker

The active platform termed “the shaker” was already introduced in ^11^. The shaker can generate single or combined horizontal, vertical, and tilting perturbations with a payload up to 1 kg. To make it suitable for small animals during striding locomotion, the shaker was conceived to generate horizontal and vertical perturbations with amplitudes of up to 1 cm at oscillation frequencies of up to 10 Hz (repeat accuracy < 0.1 mm). Tilting perturbations are only used in posture experiments and therefore do not need to satisfy such high oscillation rates. The support platform consists of two carbon plates with an elastomer in between (thickness = 6 mm) and two or three carbon tubes (external diameter ϕ = 25 mm, wall thickness = 1 mm). On the support platform, a ring made of Polyoxymethylene (POM) was rigged to permit the mounting of up to four force plates based on ATI-Nano 17® force/torque sensors ^51^. Four plates made of acrylic glass (10 cm x10 cm x 0.5 cm) were used as a platform for collecting GRFs and center of pressure (CoP) (see Fig. 6-left). Due to restrictions in the data acquisition software and a reduced x-ray window (see below), we only instrumentalized the left part of the shaker. The two plates on the right side of the shaker have “dummy” sensors made of aluminum.

**Figure 6:**
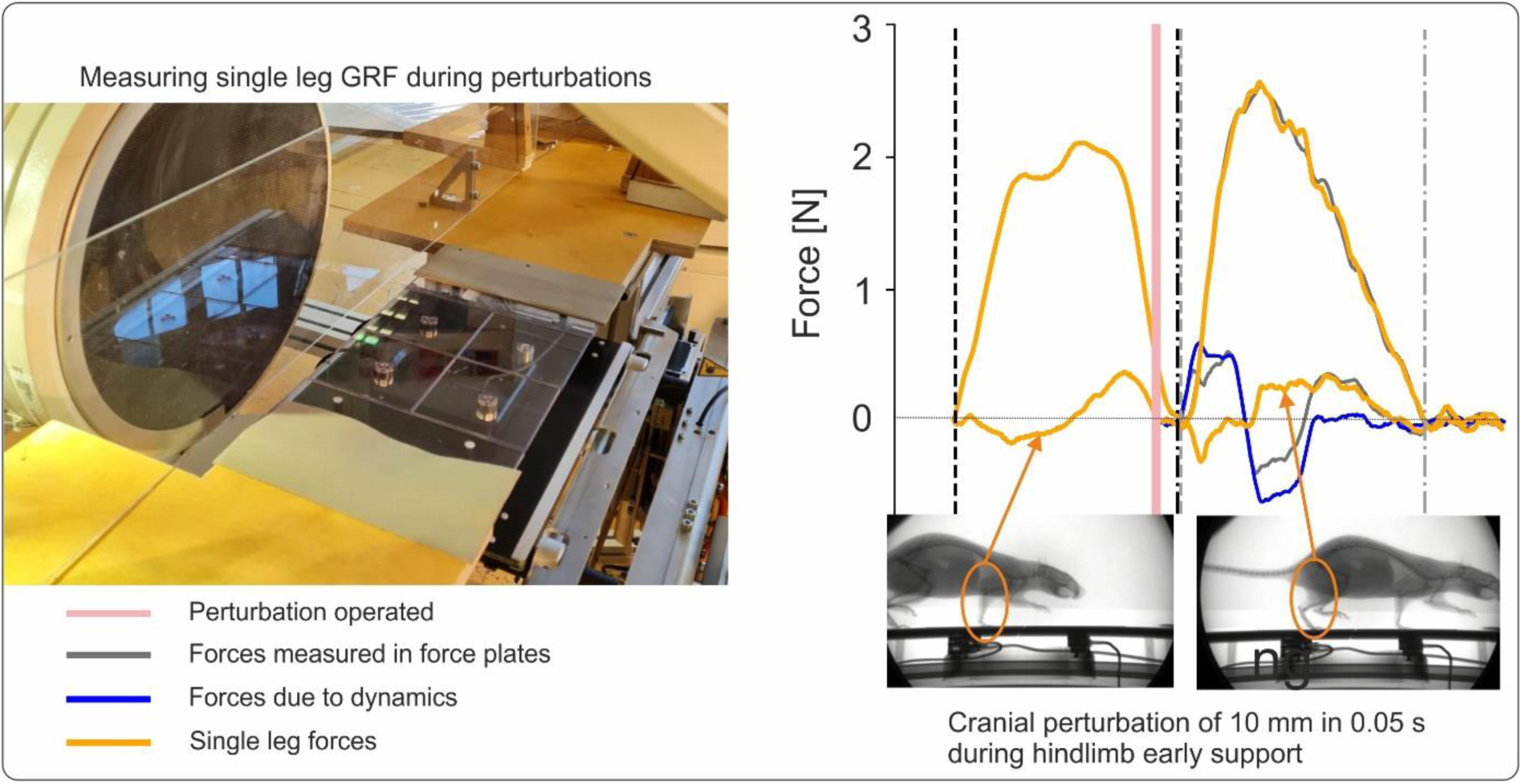
Analyzing actively perturbed locomotion in rats. Left) Experimental setup combining biplanar X-ray and an active platform (“shaker”) capable of force measure. Right) ground reaction forces before (right forelimb) and during a horizontal perturbation (right hindlimb). The platform was shifted 10 mm in 0.05 s in a cranial direction at the beginning of the hindlimb support phase. Grey lines represent the forces measured by the plates, blue lines the forces produced by the dynamics of the motion (previously saved), and orange the forces exerted by the rat.

**Figure 7.**
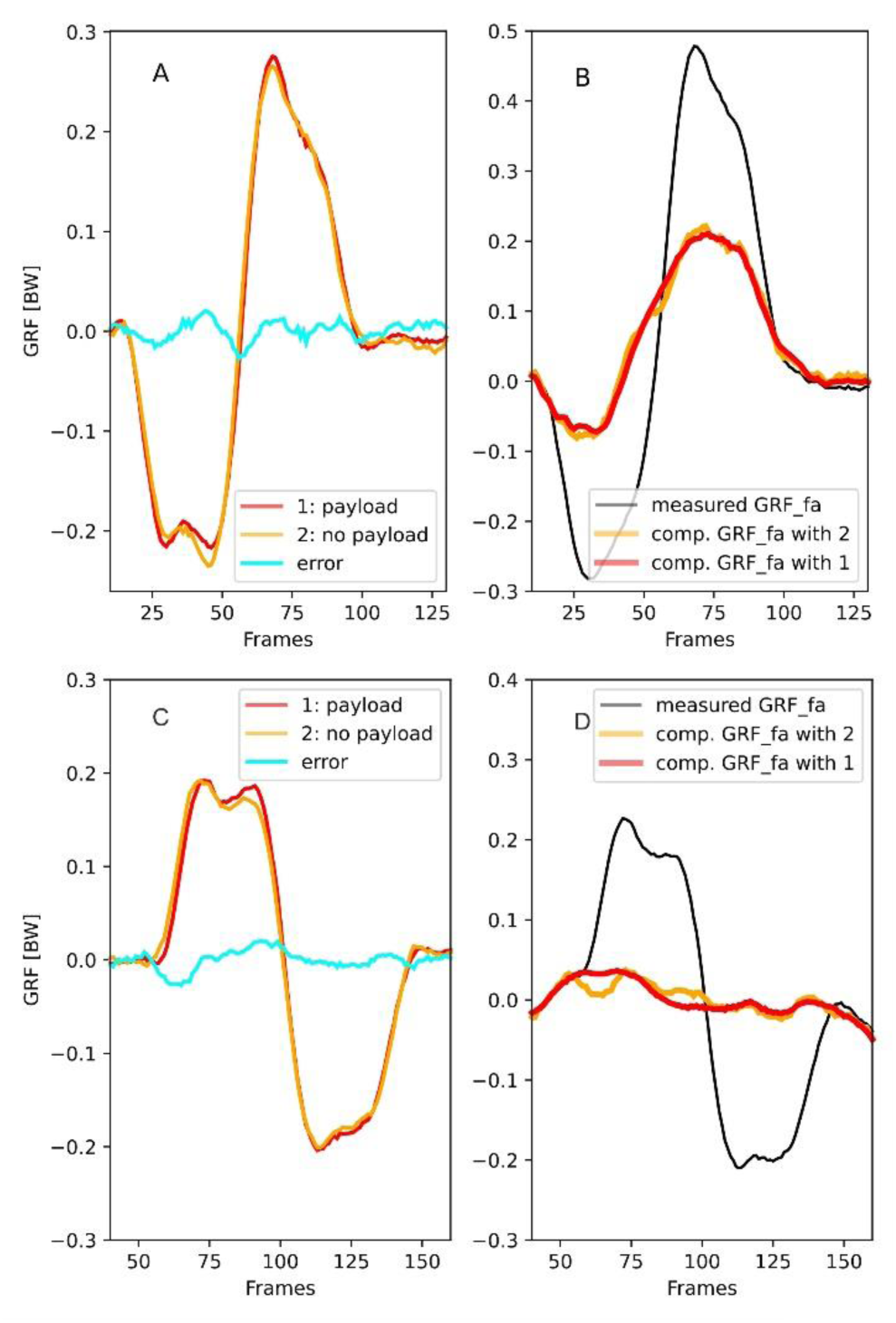
Error in the estimation of the GRFfa obtained by subtracting the force measured by the force plates without payload (orange) from those obtained with the rat on the platform (red). A-B: Caudal perturbation of 10mm in 0.05s, Cau10, C-D: Cranial perturbation of 10mm in 0.05 s. In these trials, the rats (“Red_1” in A – B, and “Right” in C-D) missed one force plate during the perturbation. Thus, that force plate measured only the forces generated by the motion of the shaker (see A & C, red curves). For these trials, it is possible to compute the error (A & C, cyan curves) introduced when using the forces obtained from the displacement of the shaker without payload (A & C, orange curves) to estimate the GRFfa. The right panel displays the effect of using both compensation forces on the measured GRFfa. Error can produce small bumps/dips or oscillations of maximum 0.03 BW.

The vertical oscillations of the platform (z-axis) are produced by two linear servomotors (Beckhoff, mod. AL8042). The horizontal oscillations (x-axis) are produced by one linear servomotor (Beckhoff, mod. AL8040). The linear motors are controlled by digital compact servo drives (Beckhoff, mod. AX5206 /Beckhoff, mod AX5106). The maximal translation amplitude ratio is 2 cm in 0.05 s. GUI and control programs were written in Beckhoff’s own software environment (Beckhoff TwinCAT). Time-dependent perturbation profiles can be freely designed using coma separated values (csv. file). Perturbations can be triggered manually or by means of a photoelectric sensor. The shaker was constructed by H&S Robotics, Ilmenau, Germany.

### Animals, experiments, and data analyses

Ten adult rats (Rattus Norvegicus) weighing from 250 g to 400 g (see Table 1) moved across a 2.3 m walking track constructed around the shaker. The Committee for Animal Research of the State of Thuringia, Germany, approved the animal care and all experimental procedures (registry number: 02-060/16).

In our experiments, the rats trotted at self-selected speeds. In the first set of experiments, several unperturbed trials were carried out to build up a basis for comparison. Note that rats, like many small prey animals, do not necessarily move steadily. For this reason, we favored the term unperturbed locomotion over steady-state locomotion. In a second set of experiments, the rats randomly experienced: a) no perturbations, or b) one-way linear perturbations of 5 mm or 10 mm in 0.05 s when they stepped on the shaker. Perturbation onset was manually triggered by the experimenter. Perturbations were vertical (up/ down) or horizontal (forth/ back). Here we present results from horizontal perturbations. Horizontal perturbations were termed cranial or caudal based on the direction of motion of the platform relative to the animal (see Fig. 1). Cranial perturbation refers to the forward motion of the platform while caudal perturbation describes the backwards motion of the platform. The duration of perturbation (0.05s) was chosen to make up about 50% of the stance time during the fast trot in rats. 0.05s is about five times the conduction delay (time required by a limb to respond to a perturbation) in a rat ^38^. Furthermore, the chosen perturbation time is also long enough to take account of other sources of delay in responses to stimuli, such as synaptic transmission and force generation^38,52^. In other words, rats in our experiments had enough time during/after perturbations for neural feedback^10^ including compensation through internal models ^53^.

Body and limb kinematics were collected using a biplanar high-speed X-ray fluoroscope (Neurostar, Siemens, Erlangen, Germany) and two synchronized standard live high-speed cameras (SpeedCam Visario g2, Weinberger, Erlangen, Germany). One plane of the X-ray machine recorded the motions of the rat in the sagittal plane (see Fig. 6-left). The second plane, which normally records from above the animal, was rotated 30° from the vertical position to minimize interference with the force/torque sensors and to improve recognition of the tantalum beads after motion capture. The X-ray machine parameters were 55 kV and 40 mA, with a sampling frequency of 500 Hz. GRFs were collected at 1.0 kHz and force and X-ray data synchronized electronically (post-trigger). Force data were synchronized with the onset of perturbation using a visual signal integrated into the shaker which was captured by the live high-speed cameras. This synchronization was necessary to permit a subtraction of the forces induced by acceleration (measured without payload) from those collected during posture/locomotion experiments (see Fig. 6-right). Note that the motion profiles produced by the linear servomotor (Beckhoff, mod. AL8040) were similar whether the rat was on the platform or off it (Fig. 7 A-C). Therefore, subtracting the forces obtained without payload from those obtained during actual trials returns a satisfactory approximation of the leg load during perturbations. However, compensating for forces obtained during motion without payload can introduce small bumps or oscillations (we measured a maximum amplitude of 0.03 BW) in the force data (see Fig. 7 B-C). The timepoint at which errors are added to the force data depends on the accuracy of the shaker’s motion and the synchronization error.

Rough force data was filtered using a 7th-order Butterworth low-pass filter with a cut-off frequency of 100 Hz applied in a zero-phase digital filter. Prior to analysis, filtered data was interpolated to 100 points between TD and TO. Double support phases occurring on the same force plate were deleted from the interpolated force data. Results were grouped and then exported to cvs-tables based on perturbation type (cra5: cranial perturbation of 5mm in 0.05s, cra10: cranial perturbation of 10mm in 0.05s, cau5: caudal perturbation of 5mm in 0.05s, cau10: caudal perturbation of 10mm in 0.05s), GRF components (vertical, fore-aft), speed group based on contact time (CT) (moderate trot CT > 110 ms, vs. fast trot CT<= 110 ms), and onset of the perturbation (early stance phase: from 0% to 50% and late stance phase: 51% to 99%). While the speed groups might reduce the effects of speed on the results, the timepoint of onset of the perturbation might roughly reflect the effects of leg orientation and perhaps muscle state in the response to perturbations.

### Statistical analysis

We analyzed CTs and GRFs at several points. Subsets of data were created for vertical and fore-aft GRFs containing information about minimum and/or maximum values and the timepoint of their occurrence. Those data subsets and CT data were afterwards split into five perturbation groups: unperturbed, Cau5, Cau10, Cra5, Cra10, and two speed categories: moderate and fast trots.

Each of these data subsets was then analyzed separately for the fore and hind limb and for early or late stance perturbations. Analysis of variance (repeated measures Anova) was performed to test whether the perturbation groups differed significantly (R:aov; ^54^). Based on this, a post-hoc test (TukeyHSD) was performed to compare all possible groups ^55^. As homogeneous variance could not be assumed for all parameters, a variance test (Levene) was also carried out. In the case of problematic variances, all possible groups were compared using a further post-hoc test (Games-Howell; ^56^). The statistical analysis was carried out using the freely available software R (version: 4.2.3). The libraries (R.matlab, data.table, Rfast, tidyverse, stats, rstatix, car, ggstatsplot, viridis, openxlsx) were used for specific functions. R-scripts were created using the free program “master” (https://starkrats.de).

Because statistical analyses mostly failed to detect differences between unperturbed and perturbed locomotion, we introduced a measure that we termed impulse gap (IG) to classify perturbed trials relative to unperturbed ones. IG provides the sum of the distances (the area) between the curve representing the mean value of GRF_fa_ obtained during unperturbed trials (mean GRF_faUnpert_) and the GRF_fa_ curve of a perturbed trial (GRF_faPert_) for *i* frames. Thus *IG* = ∑^*n*^_i=0_ (*mean GRF*_*faUnpert_i_*_ − *GRF*_*faPert_i_*_). In other words, IG is a measure of the shift between perturbed and unperturbed fore-aft GRF during the analyzed period. We then compared IG with the impulse margins (ISD) obtained during unperturbed locomotion. ISD represents the area computed between the mean GRF_fa_ for unperturbed trials (GRF_faUnpert_) and the standard deviation (SD) of GRF_faUnpert_, *ISD* = ∑^*n*^ (*mean GRF*_*faUnpert_i_*_ − *SD GRF*_*faUnpert_i_*_). If IG > ISD, the leg accelerated during the step/part of the step relative to the averaged unperturbed step. Conversely, if IG < -ISD, the leg braked. Finally, if -ISD < IG < ISD, it means that the impulse gap remained within the impulse value measured between mean GRF_faUnpert_ ± SD GRF_faUnpert_. This is a necessary condition to categorize a step as a non-resetting step. However, a GRF_faPert_ that oscillates around the mean GRF_faUnpert_ could even with larger amplitudes return very low IG values. Therefore, IG^2^ must also be lower than or equal to ISD^2^ in order to be able to categorize a step as non-resetting. We analyzed IG, IG^2^, ISD and ISD^2^ for the whole step, and for the portions of the step before and after the perturbation. Measuring IG for the whole step provides information about the general behavior during the contact phase. Measuring IG before the perturbation may provide information about feedforward strategies, while after the perturbation it provides an insight into the behavioral reaction to the perturbation. For every behavior, we then inspected individual trials to look for perturbation-related alterations to the shape of the GRF curve after the triggering of the horizontal shifts. Data analysis was performed using Python (versions 3.9.7 and 3.11.5) and the following Python libraries: Pandas, Scipy, Numpy, Pickle. Plots were produced using Matplotlib.pyplot.

## Declarations

### Ethics approval and consent to participate

All experiments were approved by and carried out in strict accordance with the German Animal Welfare guidelines and regulations of the states of Thuringia (registry number: 02-060/16). We confirm that we complied with the ARRIVE guidelines.

### Consent for publication

Not applicable

### Availability of data and materials

The datasets used and/or analyzed during the current study are available from the corresponding author on reasonable request.

### Authors’ contributions

E.A., D.A., and M.S.F. designed the study. E.A., D.A., and M.S.F. supervised and performed the experiments. E.A. analyzed experimental data, H.S. performed the statistical analysis. E.A. and M.S.F. were responsible for grant acquisition. E.A. drafted the manuscript and all figures. All authors contributed to the interpretation of the results and revised the manuscript.

## Acknowledgements

We thank Matthew C. Tresch for valuable feedback and Lucy Cathrow language polishing. We also thank Ingrid Weiss and Rommy Petersohn for their technical assistance during the experiments.

## List of abbreviations

BW: Force expressed in body weight

Cau5: 5 mm caudal pertubation

Cau10: 10 mm caudal pertubation

Cra5: 5 mm cranial pertubation

Cra10: 10 mm cranial pertubation

CT: Contact time

ESP: Early stance perturbation

GRF: Ground reaction forces

GRF_v_: Vertical component of the ground reaction forces

GRF_fa_: Fore-aft component of the ground reaction forces

IG: Impulse gap

ISD: Impulse margin for unperturbed locomotion

LSP: Late stance perturbation

SD: standard deviation

SLGRF: Single leg ground reaction forces

TD: Touch-down

TO: Toe-off

## References

1 Blickhan, R. et al. Intelligence by mechanics. Philos Transact A Math Phys Eng Sci 365, 199–220, doi:10.1098/rsta.2006.1911 (2007).

2 Dickinson, M. H. et al. How animals move: an integrative view. Science 288, 100–106 (2000).

3 Nishikawa, K. et al. Neuromechanics: an integrative approach for understanding motor control. Integrative and Comparative Biology 47, 16–54, doi:10.1093/icb/icm024 (2007).

4 Biewener, A. A. & Daley, M. A. Unsteady locomotion: integrating muscle function with whole body dynamics and neuromuscular control. Journal of Experimental Biology 210, 2949–2960, doi:10.1242/jeb.005801 (2007).

5 Birn-Jeffery, A. V. & Daley, M. A. Birds achieve high robustness in uneven terrain through active control of landing conditions. The Journal of Experimental Biology 215, 2117–2127, doi:10.1242/jeb.065557 (2012).

6 Birn-Jeffery, A. V. et al. Don’t break a leg: running birds from quail to ostrich prioritise leg safety and economy on uneven terrain. Journal of Experimental Biology 217, 3786–3796 (2014).

7 Daley, M. A. & Biewener, A. A. Leg muscles that mediate stability: mechanics and control of two distal extensor muscles during obstacle negotiation in the guinea fowl. Philosophical Transactions of the Royal Society B: Biological Sciences 366, 1580–1591 (2011).

8 Gordon, J. C., Rankin, J. W. & Daley, M. A. How do treadmill speed and terrain visibility influence neuromuscular control of guinea fowl locomotion? Journal of Experimental Biology 218, 3010–3022 (2015).

9 Andrada, E. et al. Limb, joint and pelvic kinematic control in the quail coping with steps upwards and downwards. Scientific reports 12, 1–17 (2022).

10 Sponberg, S., Abbott, E. & Sawicki, G. S. Perturbing the muscle work loop paradigm to unravel the neuromechanics of unsteady locomotion. Journal of Experimental Biology 226, jeb243561 (2023).

11 Andrada, E., Karguth, A. & Fischer, M. S. in Biomimetic and Biohybrid Systems. (eds Alexander Hunt et al.) 103-106 (Springer International Publishing).

12 Muir, G. D. & Webb, A. A. Assessment of behavioural recovery following spinal cord injury in rats. European Journal of Neuroscience 12, 3079–3086 (2000).

13 Muir, G. D. & Whishaw, I. Q. Complete locomotor recovery following corticospinal tract lesions: measurement of ground reaction forces during overground locomotion in rats. Behavioural brain research 103, 45–53 (1999).

14 Muir, G. D. & Whishaw, I. Q. Ground reaction forces in locomoting hemi-parkinsonian rats: a definitive test for impairments and compensations. Experimental Brain Research 126, 307–314 (1999).

15 Heglund, N. C. A simple design for a force-plate to measure ground reaction forces. (1981).

16 Biewener, A. A. Animal locomotion. (Oxford University Press, 2003).

17 Clarke, K. Differential fore-and hindpaw force transmission in the walking rat. Physiology & behavior 58, 415–419 (1995).

18 Andrada, E., Mämpel, J., Schmidt, A., Fischer, M. S. & Witte, H. From Biomechanics Of Rats’ Inclined Locomotion To A Climbing Robot. International Journal of Design & Nature and Ecodynamics 8, 191–212 (2013).

19 Wildau, J., Arnold, D., Rode, C., Andrada, E. & Fischer, M. in JOURNAL OF MORPHOLOGY. S238-S238 (WILEY 111 RIVER ST, HOBOKEN 07030-5774, NJ USA).

20 Johnson, W. L., Jindrich, D. L., Roy, R. R. & Edgerton, V. R. Quantitative metrics of spinal cord injury recovery in the rat using motion capture, electromyography and ground reaction force measurement. Journal of Neuroscience Methods 206, 65–72 (2012).

21 Brandt, T., Strupp, M. & Benson, J. You are better off running than walking with acute vestibulopathy. The Lancet 354, 746 (1999).

22 Mergner, T. & Rosemeier, T. Interaction of vestibular, somatosensory and visual signals for postural control and motion perception under terrestrial and microgravity conditions—a conceptual model. Brain research reviews 28, 118–135 (1998).

23 Dietz, V. Human neuronal control of automatic functional movements: interaction between central programs and afferent input. Physiological reviews 72, 33–69 (1992).

24 Grillner, S. & Rossignol, S. On the initiation of the swing phase of locomotion in chronic spinal cats. Brain research 146, 269–277 (1978).

25 Whelan, P. J. Control of locomotion in the decerebrate cat. Progress in neurobiology 49, 481–515 (1996).

26 Kiehn, O. Decoding the organization of spinal circuits that control locomotion. Nature Reviews Neuroscience 17, 224–238 (2016).

27 Grillner, S. & Kozlov, A. The CPGs for limbed locomotion–facts and fiction. International Journal of Molecular Sciences 22, 5882 (2021).

28 Grillner, S. & Manira, A. E. Current Principles of Motor Control, with Special Reference to Vertebrate Locomotion. Physiological Reviews 100, 271–320, doi:10.1152/physrev.00015.2019 (2020).

29 Marigold, D. S. & Patla, A. E. Adapting Locomotion to Different Surface Compliances: Neuromuscular Responses and Changes in Movement Dynamics. Journal of Neurophysiology 94, 1733–1750, doi:10.1152/jn.00019.2005 (2005).

30 Mohagheghi, A. A., Moraes, R. & Patla, A. E. The effects of distant and on-line visual information on the control of approach phase and step over an obstacle during locomotion. Experimental Brain Research 155, 459–468 (2004).

31 Patla, A. E. How Is Human Gait Controlled by Vision. Ecological Psychology 10, 287–302, doi:10.1080/10407413.1998.9652686 (1998).

32 Rybak, I. A., Stecina, K., Shevtsova, N. A. & McCrea, D. A. Modelling spinal circuitry involved in locomotor pattern generation: insights from the effects of afferent stimulation. The Journal of physiology 577, 641–658 (2006).

33 Granatosky, M. C. et al. Variation in limb loading magnitude and timing in tetrapods. The Journal of Experimental Biology 223, jeb201525, doi:10.1242/jeb.201525 (2020).

34 Aoi, S. et al. A stability-based mechanism for hysteresis in the walk–trot transition in quadruped locomotion. Journal of The Royal Society Interface 10, 20120908 (2013).

35 Ross, C. F. et al. The evolution of locomotor rhythmicity in tetrapods. Evolution 67, 1209–1217 (2013).

36 Hulliger, M. The mammalian muscle spindle and its central control. *Reviews of Physiology, Biochemistry and Pharmacology*, Volume 101: *Volume: 101*, 1-110 (1984).

37 McMahon, T. A. Muscles, reflexes, and locomotion. Vol. 10 (Princeton University Press, 1984).

38 More, H. L. et al. Scaling of sensorimotor control in terrestrial mammals. Proceedings of the Royal Society B: Biological Sciences 277, 3563–3568 (2010).

39 Grottel, K. & Celichowski, J. Division of motor units in medial gastrocnemius muscle of the rat in the light of variability of their principal properties. Acta Neurobiologiae Experimentalis 50, 571–587 (1990).

40 Fischer, M. S. & Blickhan, R. The Tri-Segmented Limbs of Therian Mammals: Kinematics, Dynamics, and Self-stabilization-A Review. J. Exp. Zool., 935–952 (2006).

41 Daley, M. A., Voloshina, A. & Biewener, A. A. The role of intrinsic muscle mechanics in the neuromuscular control of stable running in the guinea fowl. The Journal of physiology 587, 2693–2707 (2009).

42 Josephson, R. K. Dissecting muscle power output. Journal of Experimental Biology 202, 3369–3375, doi:10.1242/jeb.202.23.3369 (1999).

43 Taylor, A., Durbaba, R., Ellaway, P. & Rawlinson, S. Static and dynamic γ-motor output to ankle flexor muscles during locomotion in the decerebrate cat. The Journal of physiology 571, 711–723 (2006).

44 Taylor, A., Ellaway, P., Durbaba, R. & Rawlinson, S. Distinctive patterns of static and dynamic gamma motor activity during locomotion in the decerebrate cat. The Journal of physiology 529, 825–836 (2000).

45 Caicoya, A., Illert, M. & Jänike, R. Monosynaptic Ia pathways at the cat shoulder. The Journal of Physiology 518, 825–841 (1999).

46 Marigold, D. S. & Patla, A. E. Strategies for Dynamic Stability During Locomotion on a Slippery Surface: Effects of Prior Experience and Knowledge. J of Neurophysiol 88, 339–353, doi:10.1152/jn.00691.2001 (2002).

47 Pearson, K. Motor systems. Current opinion in neurobiology 10, 649–654 (2000).

48 Russell, D. F. & Zajac, F. Effects of stimulalating Deiters’ nucleus and memdial longitudinal fasciculus on the timing of the fictive locomotor rhythm induced in cats by DOPA. Brain Research 177, 588–592, 10.1016/0006-8993(79)90478-5 (1979).

49 Wilson, H. R. Spikes, decisions, and actions: the dynamical foundations of neuroscience. (No Title) (1999).

50 Sutton, G., Szczecinski, N., Quinn, R. & Chiel, H. Neural control of rhythmic limb motion is shaped by size and speed. (2021).

51 Andrada, E., Nyakatura, J. A., Bergmann, F. & Blickhan, R. Adjustments of global and local hindlimb properties during terrestrial locomotion of the common quail (Coturnix coturnix). The Journal of Experimental Biology 216, 3906–3916 (2013).

52 Enoka, R. M. Morphological features and activation patterns of motor units. Journal of Clinical Neurophysiology 12, 538–559 (1995).

53 Wolpert, D. M. & Ghahramani, Z. Computational principles of movement neuroscience. Nature neuroscience 3, 1212–1217 (2000).

54 Chambers, J. M., Freeny, A. E. & Heiberger, R. M. in Statistical models in S 145–193 (Routledge, 2017).

55 Yandell, B. Practical data analysis for designed experiments. (Routledge, 2017).

56 Ruxton, G. D. & Beauchamp, G. Time for some a priori thinking about post hoc testing. Behavioral ecology 19, 690–693 (2008).

