## Supplementary Tables for "Control of limb loading during active horizontal perturbations at moderate and fast trots in rats"

The complete set of statistical tests and their results can be found under following link: http://rat.stark-jena.de.

**Table S2. Contact times [ms] for unperturbed and early stance perturbations.**

Forelimbs

| Perturbation | n | min | max | median | mean | SD |
|---|---|---|---|---|---|---|
| Unpert. (mod) | 31 | 110 | 292 | 160 | 166,2 | 40,8 |
| Unpert. (fast) | 10 | 70 | 102 | 88 | 88,2 | 9,18 |
| Cau5 (mod) | 11 | 114 | 246 | 135 | 148,5 | 39,3 |
| Cau5 (fast) | 3 | 82 | 92 | 86 | 86, 7 | 5,0 |
| Cau10 (mod) | 7 | 120 | 167 | 148 | 147,7 | 16,4 |
| Cau10 (fast) | 5 | 66 | 108 | 100 | 94,6 | 16,5 |
| Cra5 (mod) | 19 | 110 | 232 | 176 | 170,9 | 38,9 |
| Cra5 (fast) | 3 | 76 | 106 | 76 | 86 | 17,3 |
| Cra10 (mod) | 12 | 112 | 212 | 149 | 146,8 | 31,4 |
| Cra10 (fast) | 2 | 76 | 108 | 92 | 92 | 22,6 |

Hindlimbs

| Perturbation | n | min | max | median | mean | SD |
|---|---|---|---|---|---|---|
| Unpert. (mod) | 26 | 111 | 286 | 166 | 164 | 37,7 |
| Unpert. (fast) | 10 | 66 | 104 | 90 | 87 | 13,6 |
| Cau5 (mod) | 12 | 116 | 192 | 146 | 152 | 30,8 |
| Cau5 (fast) | 5 | 74 | 106 | 102 | 94 | 13,9 |
| Cau10 (mod) | 8 | 132 | 182 | 150 | 155 | 19,7 |
| Cau10 (fast) | 7 | 78 | 106 | 102 | 96 | 12,4 |
| Cra5 (mod) | 16 | 112 | 242 | 151,5 | 156 | 36,5 |
| Cra5 (fast) | 4 | 68 | 107 | 97 | 92 | 17,0 |
| Cra10 (mod) | 15 | 110 | 254 | 148 | 155 | 44,1 |
| Cra10 (fast) | 4 | 94 | 106 | 103 | 102 | 5,3 |

SD: Standard deviation, n: Number of steps, mod: Moderate trot, fast: Fast trot.

**Table S3. Contact times [ms] for late stance perturbations.**

| | | | Forelimbs | | | |
|---|---|---|---|---|---|---|
| Perturbation | n | min | max | median | mean | SD |
| Cau5 (mod) | 12 | 112 | 224 | 153 | 154,6 | 33,4 |
| Cau5 (fast) | 6 | 78 | 106 | 90 | 91,3 | 11,2 |
| Cau10 (mod) | 12 | 112 | 270 | 156 | 161 | 43,6 |
| Cau10 (fast) | 3 | 98 | 108 | 98 | 101,3 | 5,8 |
| Cra5 (mod) | 19 | 114 | 249 | 140 | 152,8 | 35,8 |
| Cra5 (fast) | 4 | 84 | 96 | 94 | 92 | 5,7 |
| Cra10 (mod) | 12 | 112 | 258 | 159 | 175 | 45, 8 |
| Cra10 (fast) | 2 | 80 | 90 | 85 | 85 | 7,1 |
| | | | Hindlimbs | | | |
| Perturbation | n | min | max | median | mean | SD |
| Cau5 (mod) | 5 | 132 | 256 | 178 | 188 | 45,6 |
| Cau5 (fast) | 5 | 72 | 106 | 90 | 91,6 | 13,4 |
| Cau10 (mod) | | | | | | |
| Cau10 (fast) | 3 | 116 | 184 | 178 | 159,3 | 37,6 |
| Cra5 (mod) | 5 | 138 | 255 | 174 | 181 | 44,7 |
| Cra5 (fast) | 3 | 90 | 108 | 92 | 96, 7 | 9,9 |
| Cra10 (mod) | 5 | 128 | 172 | 164 | 154,8 | 18,3 |
| Cra10 (fast) | 2 | 82 | 108 | 95 | 95 | 18,4 |

SD: Standard deviation, n: Number of steps, mod: Moderate trot, fast: Fast trot.

**Table S4. Braking fore-aft forces, min values [BW] during early stance perturbations.**

| Perturbation | n | min | max | median | mean | SD |
|---|---|---|---|---|---|---|
| | | | Forelimbs | | | |
| Perturbation | n | min | max | median | mean | SD |
| Unpert. (mod) | 31 | -0,18 | -0,03 | -0,10 | -0,10 | 0,04 |
| Unpert. (fast) | 10 | -0,13 | -0,03 | -0,08 | -0,09 | 0,03 |
| Cau5 (mod) | 11 | -0,13 | -0,015 | -0,07 | -0,07 | 0,04 |
| Cau5 (fast) | 3 | -0,25 | -0,09 | -0,10 | -0,15 | 0,09 |
| Cau10 (mod) | 7 | -0,24 | -0,03 | -0,13 | -0,11 | 0,07 |
| Cau10 (fast) | 5 | -0,20 | -0,03 | -0,12 | -0,11 | 0,07 |
| Cra5 (mod) | 19 | -0,26 | -0,06 | -0,1 | -0,13 | 0,07 |
| Cra5 (fast) | 3 | -0,19 | -0,11 | -0,11 | -0,14 | 0,05 |
| Cra10 (mod) | 12 | -0,14 | -0,02 | -0,08 | -0,08 | 0,04 |
| Cranial 10 (fast) | 2 | -0,19 | -0,06 | -0,13 | -0,13 | 0,1 |
| | | | Hindlimbs | | | |
| Perturbation | n | min | max | median | mean | SD |
| Unpert. (mod) | 25 | -0,18 | -0,03 | -0,091 | -0,09 | 0,03 |
| Unpert. (fast) | 10 | -0,12 | -0,04 | -0,084 | -0,08 | 0,03 |
| Cau5 (mod) | 12 | -0,16 | -0,01 | -0,10 | -0,1 | 0,05 |
| Cau5 (fast) | 5 | -0,21 | -0,03 | -0,06 | -0,08 | 0,07 |
| Cau10 (mod) | 8 | -0,14 | -0,09 | -0,13 | -0,12 | 0,02 |
| Cau10 (fast) | 7 | -0,19 | -0,03 | -0,06 | -0,09 | 0,06 |
| Cra5 (mod) | 16 | -0,23 | 0,002 | -0,1 | -0,1 | 0,06 |
| Cra5 (fast) | 4 | -0,11 | -0,01 | -0,06 | -0,06 | 0,04 |
| Cra10 (mod) | 15 | -0,29 | -0,001 | -0,10 | -0,11 | 0,07 |
| Cranial 10 (fast) | 4 | -0,07 | -0,01 | -0,06 | -0,05 | 0,03 |

SD: Standard deviation, n: Number of steps, mod: Moderate trot, fast: Fast trot.

**Table S5. Braking fore-aft forces, min values [BW] during late stance perturbations.**

| | | | Forelimbs | | | |
|---|---|---|---|---|---|---|
| Perturbation | n | min | max | median | mean | SD |
| Cau5 (mod) | 12 | -0,27 | -0,02 | -0,09 | -0,11 | 0,07 |
| Cau5 (fast) | 6 | -0,12 | -0,06 | -0,07 | -0,08 | 0,03 |
| Cau10 (mod) | 12 | -0,14 | -0,02 | -0,08 | -0,09 | 0,04 |
| Cau10 (fast) | 3 | -0,1 | -0,05 | -0,06 | -0,07 | 0,03 |
| Cra5 (mod) | 19 | -0,15 | -0,03 | -0,06 | -0,07 | 0,03 |
| Cra5 (fast) | 4 | -0,12 | -0,08 | -0,1 | -0,1 | 0,02 |
| Cra10 (mod) | 12 | -0,18 | -0,03 | -0,09 | -0,09 | 0,04 |
| Cranial 10 (fast) | 2 | -0,16 | -0,07 | -0,12 | -0,12 | 0,07 |
| | | | Hindlimb | | | |
| Perturbation | n | min | max | median | mean | SD |
| Cau5 (mod) | 5 | -0,14 | -0,05 | -0,11 | -0,11 | 0,04 |
| Cau5 (fast) | 5 | -0,08 | -0,02 | -0,07 | -0,06 | 0,02 |
| Cau10 (mod) | 2 | -0,13 | -0,1 | -0,11 | -0,11 | 0,02 |
| Cra5 (mod) | 4 | -0,09 | -0,01 | -0,05 | -0,05 | 0,04 |
| Cra5 (fast) | 3 | -0,09 | -0,06 | -0,07 | -0,08 | 0,01 |
| Cra10 (mod) | 4 | -0,12 | -0,03 | -0,07 | -0,07 | 0,05 |
| Cranial 10 (fast) | 2 | -0,1 | -0,06 | -0,08 | -0,08 | 0,03 |

SD: Standard deviation, n: Number of steps, mod: Moderate trot, fast: Fast trot.

**Table S6. Accelerating fore-aft forces, max values [BW] during early stance perturbations.**

| Perturbation | n | min | max | median | mean | SD |
|---|---|---|---|---|---|---|
| | | | Forelimbs | | | |
| Perturbation | n | min | max | median | mean | SD |
| Unpert. (mod) | 31 | 0,05 | 0,15 | 0,09 | 0,1 | 0,03 |
| Unpert. (fast) | 10 | 0,06 | 0,17 | 0,08 | 0,1 | 0,04 |
| Cau5 (mod) | 11 | 0,09 | 0,24 | 0,13 | 0,14 | 0,04 |
| Cau5 (fast) | 3 | 0,08 | 0,11 | 0,11 | 0,1 | 0,02 |
| Cau10 (mod) | 7 | 0,06 | 0,18 | 0,11 | 0,12 | 0,04 |
| Cau10 (fast) | 5 | 0,08 | 0,18 | 0,11 | 0,13 | 0,05 |
| Cra10 (mod) | 12 | 0,02 | 0,28 | 0,10 | 0,11 | 0,07 |
| Cranial 10 (fast) | 2 | 0,06 | 0,19 | 0,13 | 0,13 | 0,09 |
| | | | Hindlimbs | | | |
| Perturbation | n | min | max | median | mean | SD |
| Unpert. (mod) | 26 | 0,06 | 0,30 | 0,14 | 0,15 | 0,06 |
| Unpert. (fast) | 10 | 0,06 | 0,35 | 0,21 | 0,22 | 0,1 |
| Cau5 (mod) | 12 | 0,11 | 0,25 | 0,18 | 0,18 | 0,05 |
| Cau5 (fast) | 5 | 0,18 | 0,25 | 0,21 | 0,21 | 0,03 |
| Cau10 (mod) | 8 | 0,05 | 0,18 | 0,13 | 0,12 | 0,05 |
| Cau10 (fast) | 7 | 0,03 | 0,28 | 0,22 | 0,17 | 0,09 |
| Cra5 (mod) | 16 | 0,002 | 0,35 | 0,19 | 0,18 | 0,09 |
| Cra5 (fast) | 4 | 0,18 | 0,45 | 0,3 | 0,31 | 0,11 |
| Cra10 (mod) | 15 | 0,05 | 0,67 | 0,21 | 0,24 | 0,15 |
| Cranial 10 (fast) | 4 | 0,23 | 0,40 | 0,32 | 0,32 | 0,07 |

SD: Standard deviation, n: Number of steps, mod: Moderate trot, fast: Fast trot.

**Table S7. Accelerating fore-aft forces, max values [BW] during late stance perturbations.**

| | | | Forelimb | | | |
|---|---|---|---|---|---|---|
| Perturbation | n | min | max | median | mean | SD |
| Cau5 (mod) | 12 | 0,05 | 0,18 | 0,14 | 0,12 | 0,04 |
| Cau5 (fast) | 6 | 0,06 | 0,19 | 0,12 | 0,13 | 0,05 |
| Cau10 (mod) | 12 | 0,07 | 0,17 | 0,1 | 0,10 | 0,03 |
| Cau10 (fast) | 3 | 0,09 | 0,13 | 0,11 | 0,11 | 0,02 |
| Cra5 (mod) | 19 | 0,04 | 0,2 | 0,12 | 0,13 | 0,04 |
| Cra5 (fast) | 4 | 0,07 | 0,21 | 0,09 | 0,11 | 0,07 |
| Cra10 (mod) | 12 | 0,01 | 0,22 | 0,10 | 0,11 | 0,06 |
| Cranial 10 (fast) | 2 | 0,04 | 0,16 | 0,10 | 0,10 | 0,08 |
| | | | Hindlimb | | | |
| typ | n | min | max | median | mean | SD |
| Caudal 05 | 5 | 0,06 | 0,35 | 0,14 | 0,17 | 0,11 |
| Caudal 05f | 5 | 0,1 | 0,24 | 0,23 | 0,21 | 0,06 |
| Caudal 10 | 3 | 0,06 | 0,13 | 0,09 | 0,09 | 0,03 |
| Cranial 05 | 5 | 0,09 | 0,38 | 0,20 | 0,23 | 0,12 |
| Cranial 05f | 3 | 0,07 | 0,25 | 0,20 | 0,18 | 0,1 |
| Cranial 10 | 5 | 0,09 | 0,29 | 0,20 | 0,2 | 0,07 |
| Cranial 10f | 2 | 0,07 | 0,24 | 0,15 | 0,15 | 0,12 |

SD: Standard deviation, n: Number of steps, mod: Moderate trot, fast: Fast trot.

**Table S8. Vertical component of the GRFs, max values [BW] during early stance perturbations.**

| | | | Forelimbs | | | |
|---|---|---|---|---|---|---|
| Perturbation | n | min | max | median | mean | SD |
| Unpert. (mod) | 31 | 0,52 | 0,88 | 0,7 | 0,7 | 0,1 |
| Unpert. (fast) | 10 | 0,64 | 1,16 | 1,03 | 0,98 | 0,15 |
| Cau5 (mod) | 11 | 0,6 | 0,94 | 0,75 | 0,75 | 0,1 |
| Cau5 (fast) | 3 | 1,04 | 1,25 | 1,23 | 1,17 | 0,11 |
| Cau10 (mod) | 7 | 0,62 | 0,83 | 0,67 | 0,71 | 0,09 |
| Cau10 (fast) | 5 | 0,71 | 1,17 | 0,93 | 0,95 | 0,17 |
| Cra5 (mod) | 19 | 0,57 | 0,9 | 0,72 | 0,72 | 0,12 |
| Cra5 (fast) | 3 | 1,01 | 1,26 | 1,06 | 1,11 | 0,13 |
| Cra10 (mod) | 12 | 0,57 | 1,02 | 0,77 | 0,77 | 0,12 |
| Cranial 10 (fast) | 2 | 0,99 | 1,26 | 1,13 | 1,12 | 0,19 |
| | | | Hindlimbs | | | |
| Perturbation | n | min | max | median | mean | SD |
| Unpert. (mod) | 25 | 0,58 | 0,96 | 0,73 | 0,73 | 0,1 |
| Unpert. (fast) | 10 | 0,76 | 1,18 | 0,98 | 0,96 | 0,12 |
| Cau5 (mod) | 12 | 0,64 | 0,98 | 0,79 | 0,81 | 0,12 |
| Cau5 (fast) | 5 | 0,67 | 1,14 | 0,97 | 0,92 | 0,19 |
| Cau10 (mod) | 8 | 0,65 | 1,02 | 0,78 | 0,80 | 0,14 |
| Cau10 (fast) | 7 | 0,61 | 1,13 | 0,83 | 0,88 | 0,21 |
| Cra5 (mod) | 16 | 0,54 | 1,18 | 0,76 | 0,78 | 0,15 |
| Cra5 (fast) | 4 | 0,96 | 1,1 | 1,03 | 1,03 | 0,07 |
| Cra10 (mod) | 15 | 0,66 | 1,14 | 0,87 | 0,86 | 0,15 |
| Cranial 10 (fast) | 4 | 0,73 | 1,13 | 1,08 | 1,00 | 0,19 |

SD: Standard deviation, n: Number of steps, mod: Moderate trot, fast: Fast trot.

**Table S9. Vertical component of the GRFs, max values [BW] during late stance perturbations.**

| | | | Forelimb | | | |
|---|---|---|---|---|---|---|
| Perturbation | n | min | max | median | mean | SD |
| Cau5 (mod) | 12 | 0,59 | 0,97 | 0,71 | 0,75 | 0,11 |
| Cau5 (fast) | 6 | 0,93 | 1,22 | 1,08 | 1,08 | 0,10 |
| Cau10 (mod) | 12 | 0,52 | 0,90 | 0,71 | 0,72 | 0,11 |
| Cau10 (fast) | 3 | 0,73 | 0,96 | 0,87 | 0,86 | 0,12 |
| Cra5 (mod) | 19 | 0,56 | 0,93 | 0,74 | 0,74 | 0,11 |
| Cra5 (fast) | 4 | 0,99 | 1,3 | 1,09 | 1,12 | 0,15 |
| Cra10 (mod) | 12 | 0,54 | 1,21 | 0,73 | 0,77 | 0,22 |
| Cranial 10 (fast) | 2 | 1,13 | 1,19 | 1,16 | 1,16 | 0,05 |
| | | | Hindlimb | | | |
| Perturbation | n | min | max | median | mean | SD |
| Cau5 (mod) | 5 | 0,6 | 0,70 | 0,68 | 0,66 | 0,05 |
| Cau5 (fast) | 5 | 0,60 | 1,1 | 0,99 | 0,92 | 0,19 |
| Cau10 (mod) | 2 | 0,63 | 0,78 | 0,71 | 0,71 | 0,11 |
| Cra5 (mod) | 4 | 0,71 | 0,78 | 0,74 | 0,74 | 0,03 |
| Cra5 (fast) | 3 | 0,73 | 0,99 | 0,87 | 0,86 | 0,13 |
| Cra10 (mod) | 4 | 0,67 | 0,96 | 0,77 | 0,8 | 0,12 |
| Cranial 10 (fast) | 2 | 0,91 | 0,91 | 0,91 | 0,91 | 0,001 |

SD: Standard deviation, n: Number of steps, mod: Moderate trot, fast: Fast trot.

**Table S10. Behaviors computed by IG and IG$^2$ for the whole step**

| | Hindlimbs | | Forelimbs | |
|---|---|---|---|---|
| | Early (a, b, n) | Late (a, b, n) | Early (a, b, n) | Late (a, b, n) |
| CRA5 moderate | (2, 5, 6) | (2, 2, 1) | (5, 5, 4) | (8, 3, 6) |
| CRA5 fast | (1, 1, 5) | (0, 1, 2) | (4, 2, 0) | (1, 1, 4) |
| CRA10 moderate | (5, 3, 3) | (3, 0, 3) | (4, 3, 1) | (6, 3, 1) |
| CRA10 fast | (4, 1, 3) | (0, 1, 1) | (4, 1, 1) | (3, 1, 1) |
| Sub-Total | (12, 10, 17) | (5, 4, 7) | (17, 11, 6) | (18, 8, 12) |
| CAU5 mod | (4, 4, 4) | (1, 2, 2) | (8, 1, 2) | (4, 4, 4) |
| CAU5_fast | (0, 1, 4) | (0, 1, 4) | (0, 1, 2) | (3, 0, 3) |
| CAU10_mod | (0, 5, 4) | (1, 2, 1) | (4, 1, 2) | (4, 7, 1) |
| CAU10_fast | (0, 3, 4) | No data | (2, 3, 0) | (2, 1, 0) |
| Sub-Total | (4, 13, 16) | (2, 5, 7) | (14, 6, 6) | (13, 12, 7) |

IG: Impulse gap, IG$^2$: Quadratic impulse gap, a: acceleration, b: brake, n: non-resetting
